# Aggregated spatio-temporal division patterns emerge from reoccurring divisions of neural stem cells

**DOI:** 10.1101/2020.03.20.999748

**Authors:** V. Lupperger, C. Marr, P. Chapouton

## Abstract

The regulation of quiescence and cell cycle entry is pivotal for the maintenance of stem cell populations. Regulatory mechanisms however are poorly understood. In particular it is unclear how the activity of single stem cells is coordinated within the population, or if cells divide in a purely random fashion. We addressed this issue by analyzing division events in an adult neural stem cell (NSC) population of the zebrafish telencephalon. Spatial statistics and mathematical modeling of over 80,000 NSCs in 36 brains revealed weakly aggregated, non-random division patterns in space and time. Analyzing divisions at two timepoints allowed us to infer cell cycle and S-phase lengths computationally. Interestingly, we observed rapid cell cycle re-entries in roughly 15% of newly born NSCs. In agent based simulations of NSC populations, this re-dividing activity sufficed to induce aggregated spatio-temporal division patterns that matched the ones observed experimentally. In contrast, omitting re-divisions lead to a random spatio-temporal distribution of dividing cells. Spatio-temporal aggregation of dividing stem cells can thus emerge from the cell’s history, regardless of possible feedback mechanisms in the population.

## Introduction

Somatic stem and progenitor cells, basic units of tissue maintenance and growth, can be found in distinct states, either dividing or quiescent. The duration of the quiescence state is not predictable <u>(van Velthoven and Rando, 2019)</u> and its regulation has a profound impact on healthy tissue maintenance. Therefore, understanding the mechanisms of adult stem cell cycle regulation is crucial. Stem cells, when scattered and confined in areas where neighboring cells fulfill structural and niche functions, can be regulated locally, such as in the bone marrow niche <u>(Morrison and Scadden, 2014)</u>. However, in a homogeneous population of equipotent stem cells that reside under the same conditions it is unclear what drives distinct behaviors of quiescence or cell cycle entry. This raises the question of how stem cells make the decision to remain quiescent or enter the cell cycle individually and collectively.

Several determinants have been proposed as regulators of quiescence or division of stem and progenitor cells. Feedback mechanisms between stem cells and the environment have been found in several systems. In the adult mouse forebrain, neuronal activity in mossy cells or in granule cells regulates the activation of radial neural stem cells of the dentate gyrus <u>(Dong et al., 2019; Yeh et al., 2018)</u>. In the subventricular zone, neural stem cell (NSC) cycling is inhibited by direct contacts with endothelial cells <u>(Ottone et al., 2014)</u>, or by the release of miR204 from the choroid plexus <u>(Lepko et al., 2019)</u>. In mouse and zebrafish adult neurogenic zones, Notch activity maintains NSCs in quiescence <u>(Alunni et al., 2013; Chapouton et al., 2010; Ehm et al., 2010)</u>, in particular in the immediate neighborhood of dividing cells <u>(Chapouton et al., 2010)</u>. In other adult stem cell populations such as the epidermis, cell divisions are instructed by neighboring differentiating progeny <u>(Liang et al., 2017; Mesa et al., 2018)</u>. Besides signals from the environment, cell-intrinsic modulations have been shown to impact on the proliferative activity, for instance the metabolic control of lipogenesis that can induce proliferation <u>(Knobloch et al., 2013, 2017)</u>. Conversely, the expression of miR-9 <u>(Katz et al., 2016)</u> or the degradation of Ascl1 via the ubiquitin ligase Huwe1 <u>(Urbán et al., 2016)</u> are factors inducing quiescence. While a combination of signals received from the environment and intrinsic to the cells seem to influence proliferating activity, it remains to be precisely understood whether and how cells coordinate their activity with their neighbors.

Toward this end, we examined the distribution of cells in S-phase in the pallial (dorsal) neurogenic niche of the adult zebrafish telencephalon, which is located on the outer surface of the brain <u>(Folgueira et al., 2012; Mueller and Wullimann, 2009)</u>. The adult brain of the zebrafish and the telencephalon in particular are growing steadily <u>(Furlan et al., 2017)</u> but at a slower pace with increasing age <u>(Edelmann et al., 2013)</u>. Radial glia constitute the entire ventricular surface of the brain with a single layer of cell somata, extending long ramified processes through the parenchyme. They express - among others - Notch ligands, the transcription factors Her4 and Fezf2, the fatty acid binding protein BLBP, the enzyme Aromatase B, the intermediate filaments GFAP, and vimentin <u>(Berberoglu et al., 2014; Chapouton et al., 2011; März et al., 2010; Pellegrini et al., 2007; Than-Trong and Bally-Cuif, 2015)</u> and a small percentage of them is dividing at any time point of observation <u>(Chapouton et al., 2010)</u>. The behavior of radial glia in steady state or injury conditions in this area of the brain hints to their function as NSCs giving rise to intermediate dividing progenitors and directly to neurons <u>(Barbosa et al., 2015; Kroehne et al., 2011; März et al., 2010; Rothenaigner et al., 2011)</u>. We previously found that cell cycle entries in NSCs occur with aggregated spatial patterns <u>(Lupperger et al., 2018)</u>. Here, we extend our analysis to the spatio-temporal domain and analyze emerging patterns with an interdisciplinary mix of experimental and computational methods.

## Results

### Neural stem cells in S-phase reveal aggregated spatial patterns on the dorsal ventricular zone

To understand cell cycle regulation in a population of mainly quiescent stem cells, we studied the spatial distribution of cells in S-phase in the intact dorsal telencephalon (pallium) in whole-mount preparations. We used adult *gfap*:GFP transgenic zebrafish, where neural stem cells (NSCs) express EGFP <u>(Bernardos and Raymond, 2006)</u>. In order to assess which subset of cells is concomitantly in S-phase at a specific time point, we marked S-phases by the incorporation of the thymidine analogon EdU one hour before fixation, (Figure 1A) and detected them by staining in whole mount preparations (Figure 1B), followed by 3D confocal microscopy (Figure 1C). Within the set of *gfap*:GFP+ NSCs (Figure 1D), we identified the 3D coordinates of EdU+ nuclei (Figure 1E) automatically and verified them via manual inspection (Figure 1F). To analyze the emerging spatial pattern, we automatically detected the coordinates of all *gfap*:GFP+ NSCs (Figure 1G) and used them as a reference grid (see Supplementary Figure 1A). We then discriminated EdU+ cells (Figure 1H) into EdU+*gfap*:GFP+ cells, representing NSCs in S-phase (Figure 1I), and EdU+*gfap*:GFP-cells, representing intermediate progenitors in S-phase <u>(März et al., 2010)</u>. We quantified the distribution of NSCs in S-phase (Figure 1J) with an adapted discrete variant of Ripley’s K <u>(Ripley, 1976)</u> that measures the number of neighboring NSCs in S-phase observed in a particular radius, accounting for a nonrandom distribution of non-dividing NSCs and edge effects (see Methods). According to this spatial statistics measure, NSCs in S-phase reveal an aggregated pattern, which is significantly deviating from a random (Supplementary Figure 1B,C) and dispersed patterns (Supplementary Figure 1D,E) for radii > 50μm (Figure 1K). Aggregated patterns of NSCs in S-phase were reproducibly found in over 70% of all hemispheres analyzed (Supplementary Figure 1F), suggesting that S-phase entry happens in a spatially non-random manner.

**Figure 1:**
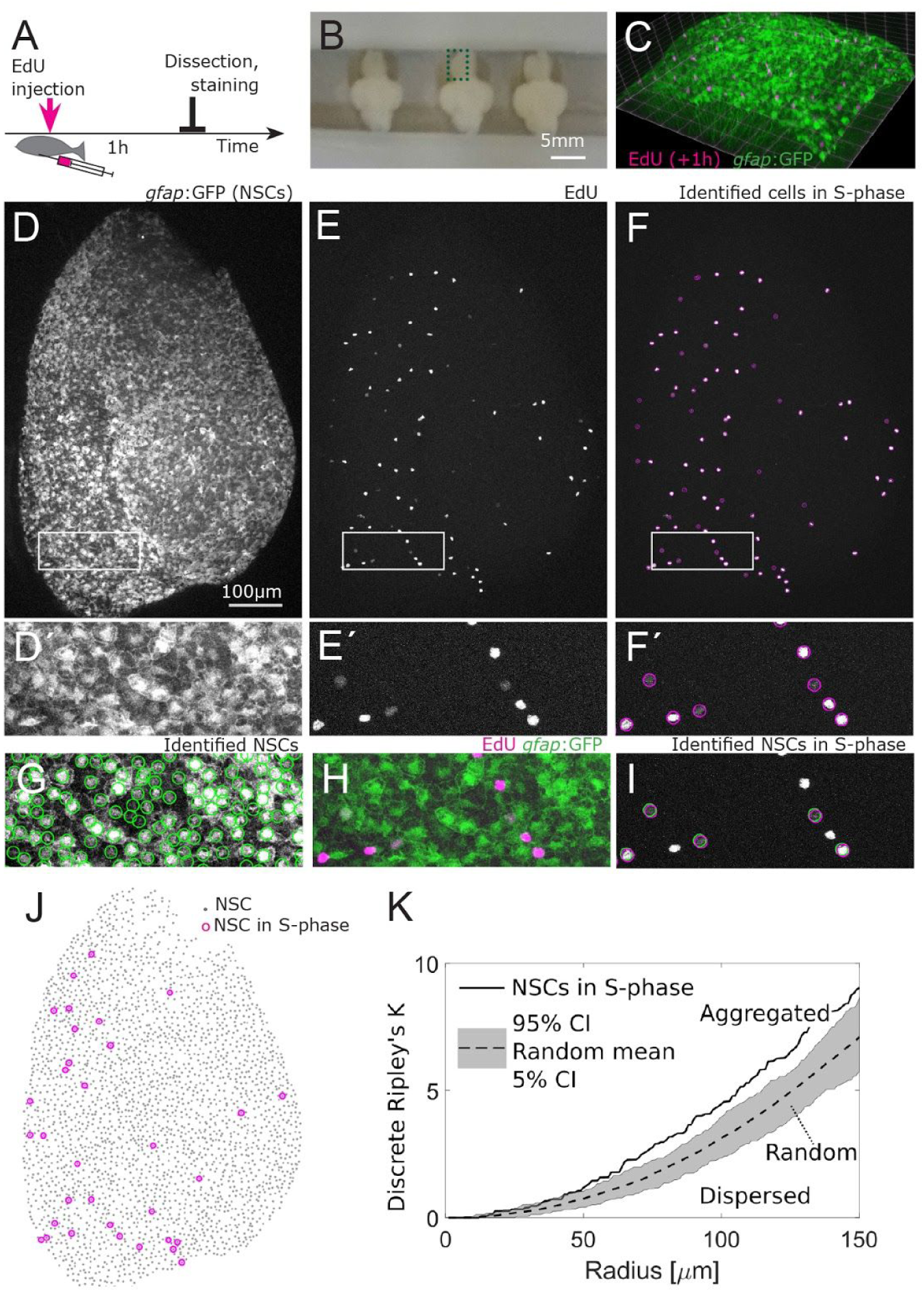
Neural stem cells (NSCs) in S-phase localize non-randomly and are spatially aggregated. (A) Experimental setup: EdU is injected intraperitoneally one hour prior to sacrificing the fish and fixation of the brain. (B) Example of three zebrafish brains mounted for 3D confocal microscopy between two coverslips separated by parafilm spacers, anterior to the top, the dorsal part is facing the bottom coverslip. The green boxed area depicts one hemisphere of the telencephalon. (C) Part of the telencephalic hemisphere as a 3D reconstruction.The *gfap*:GFP transgene highlights cell bodies of NSCs, which are arranged on a 2-dimensional layer on the telencephalon surface, while their radial processes project deep in the parenchyme. (D-D’) *Gfap*:GFP in the whole hemisphere shown as a maximum intensity z-stack projection, anterior to the top, lateral to the right, medial to the left. Boxed areas in (D-F) are shown as higher magnifications in (D’-F’). (E-E’) EdU coupled to Azyde-Alexa 647 highlights cells in S-phase and reveals their spatial distribution. (F,F’) Identified cells in S-phase surrounded by pink circles. (G) Automatically identified NSCs. (H, I) EdU+ cells were subdivided into EdU+*gfap*:GFP- and EdU+*gfap*:GFP+, the latter representing NSCs in S-phase. (J) The 33 NSCs in S-phase (pink circles) exhibit a non-random spatial pattern on top of all 2678 NSCs (gray dots). (k) Discrete Ripley’s K quantification reveals that NSCs in S-phase (solid line) are aggregated, that is, closer to each other than expected from random (dotted line with 90% confidence interval in grey) and dispersed patterns.

### Neural stem cells in S-Phase reveal aggregated spatio-temporal patterns

To investigate whether proliferative activity regulates new S-phase entries, we analyzed spatial division patterns at different time points. We made use of a second thymidine analogon, BrdU, to observe consecutive S-phases taking place *in vivo*. The two labels BrdU and EdU were administered by intraperitoneal injections separated by a labelling interval Δt of 32 hours (Figure 2A). Fish were sacrificed, their brain dissected and fixed one hour after the second administration (Figure 2A). EdU+ and BrdU+ cells were observed as clearly distinct sets (Figure 2B and 2C). When administered simultaneously as a control, EdU and BrdU reliably labelled the same set of cells (Supplementary Figure 2A). For a quantitative analysis, we identified the 3D coordinates of the BrdU+ and EdU+ NSCs (Figure 2E-G). We observed that some areas of the ventricular zone remain devoid of S-phase NSCs at those two time points. This observation was confirmed by a spatio-temporal Ripley’s K analysis: From a radius > 50 μm the density of EdU+ NSCs around BrdU+ NSCs was higher than expected from random (Figure 2H). Thus, NSCs in S-phase aggregate also spatio-temporally, a trend that was observed in all four hemispheres with a labelling interval of 32 hours (Figure 2I). These results suggest that subsequent S-phases are not randomly distributed and that the spatial organization observed is linked to the past activity in the population.

**Figure 2:**
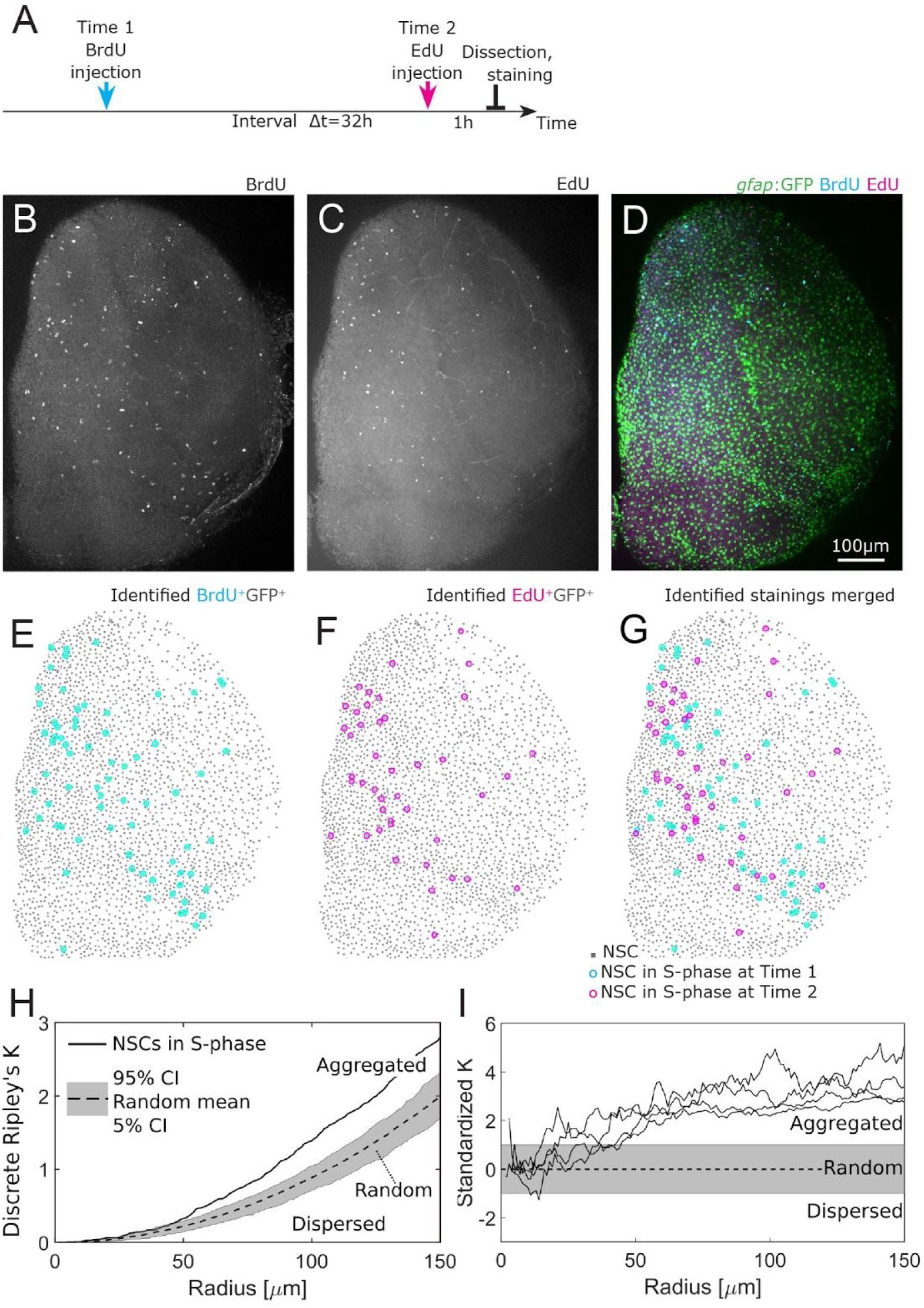
Consecutive S-phases in the NSC population are spatio-temporally aggregated. (A) Cells in S-phase are labelled with BrdU and EdU with an interval of 32h. Fish are sacrificed 1h after the EdU injection, the brains are imaged after fixation and staining. (B-D) Example of one telencephalic hemisphere, anterior to the top, lateral to the right, medial to the left, as a maximum intensity z-stack projection, in three different channels: BrdU (B), EdU (C) and *gfap*:GFP transgene highlighting NSC bodies merged with the EdU and BrdU staining (D). Scale bar: 100μm. (E-G) Identified NSCs in S-phase at two different timepoints exhibit aggregated spatio-temporal patterns. (H) Discrete Ripley’s K reveals more EdU+ NSCs around BrdU+ NSCs as expected from a random process. (I) We find spatio-temporally aggregated patterns with radii above 50μm in all 4 hemispheres where S-phases have been labelled with a labelling interval of 32h.

To measure the influence of time on the observed spatio-temporal patterns, we broadened the range of labelling intervals from 9h to 72h (Supplementary Figure 2B). Using the BrdU - EdU double labelling approach, we processed in total 36 hemispheres, and identified NSCs and NSCs in S-phase at both time points (Supplementary Table 1). Visually, the observed patterns vary from relatively homogeneous point clouds to strongly confined regions (Figure 3A). Quantitatively, we identify in 22 out of 36 hemispheres spatio-temporally aggregated patterns of divisions between two labelling time points (see Supplementary Figure 2C).

**Figure 3:**
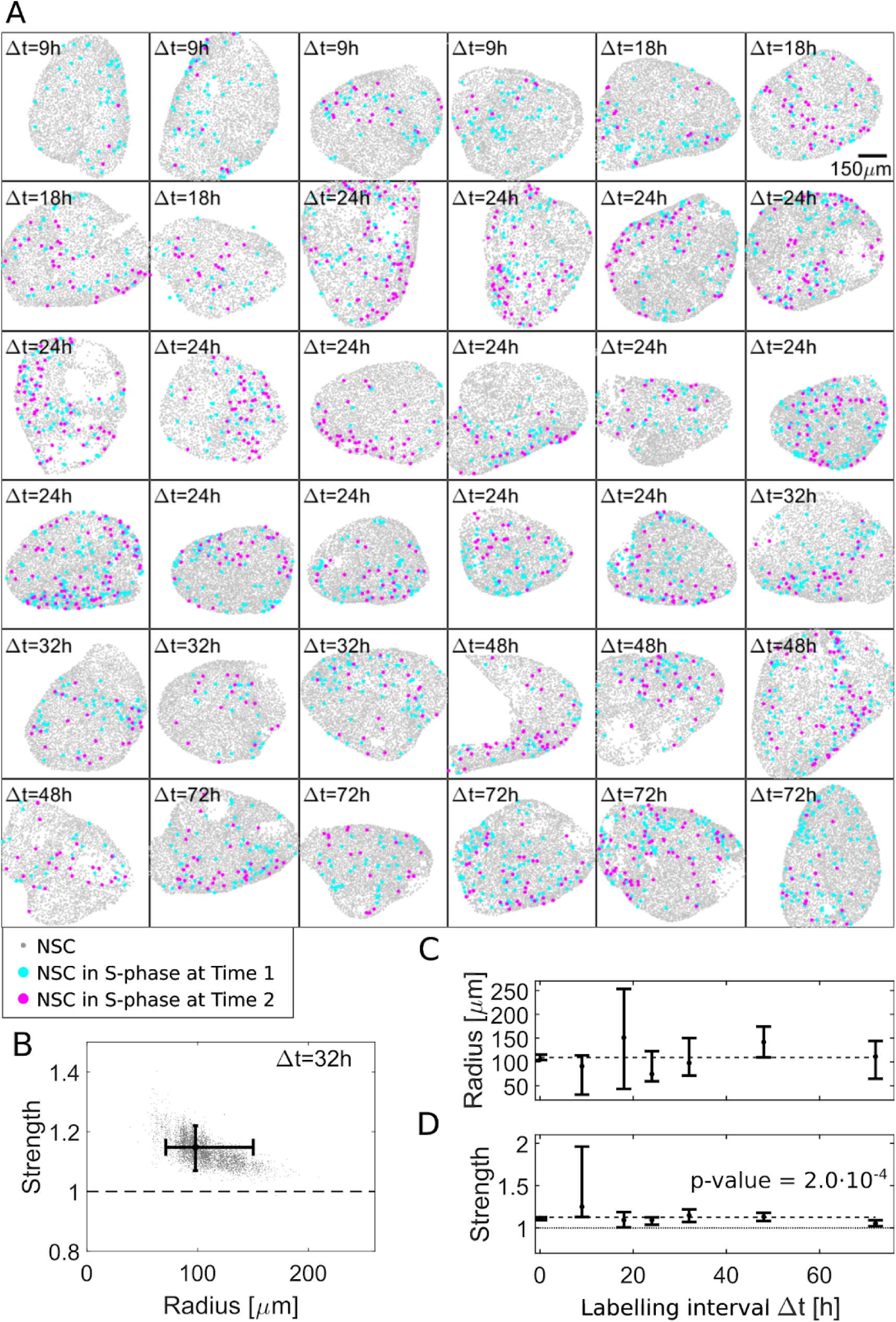
Computational approach identifies a ∼100μm aggregation radius of NSCs in S-phase. (A) Our dataset comprises 36 hemispheres with labelling intervals from Δt=9h to 72h. (B) Posterior sampling identifies the most likely strength of 1.15 and most likely radius of 98μm for four Δt=32h hemispheres. Whiskers cover the 95% confidence intervals for radius and strength. (C) Applied to all 36 hemispheres posterior sampling reveals an influence radius around 100μm. (D) The influence strength is significantly above 1 for all labelling intervals Δt (p-value = 0.0002 for constant fit to most likely values) thus inducing aggregated patterns.

### A positive interaction model fits the observed spatio-temporal patterns

Ripley’s K statistics is limited: It does not allow integrating different datasets, and it cannot quantitatively infer the strength and extent of an observed pattern. To remedy these aspects, we use the temporal extension of a spatial model <u>(Lupperger et al., 2018)</u> that allows determining the most likely parameters for interaction radius and interaction strength for an arbitrary number of datasets. Applied to the four Δt=32h patterns shown in Figure 3A, we find that a model with an interaction radius of ∼100 μm and an interaction strength > 1 describes the data best (Figure 3B, random patterns are indicated by 1, dispersed pattern by a strength < 1 and aggregated pattern by a strength > 1, see Methods). Posterior sampling reveals a 90% confidence interval from 71 to 150μm, a maximum likelihood influence radius at 98μm, and a 90% confidence interval from 1.07 to 1.22 with a maximum likelihood influence strength of 1.15. Applied to all 36 hemispheres with the coordinates of 87807 NSCs at distinct labelling intervals Δt (see Supplementary Figure 3), we find the most likely interaction radii to be around 100μm (Figure 3C). The interaction strength (Figure 3D) is robustly above 1, indicating a significant (p-value=0.0002 for a linear regression model) spatio-temporal aggregation of S-phase NSCs for all labelling intervals Δt.

### Re-division of NSC daughter cells occur within 24 to 72 hours

Our spatio-temporal analysis revealed a weak aggregation of successive NSCs in S-phase. This could arise (i) through direct stimulating contacts between dividing cells, (ii) due to local diffusible signals, or (iii) from cell-intrinsic processes driven by the cells history. We thus inquired whether a NSC’s history would be of relevance for new S-phase entries and made use of a double-labelling approach. Short labelling intervals up to Δt=24h revealed cells in S-phase straddling the Δt and incorporating both labels (Figure 4A-F), hence denoted as double labelled S-phases (DLS). Remarkably, we also noticed dividing NSCs that entered a second S-phase within labelling intervals of Δt=24h, 32h, 48h, and 72h (Figure 4G-T), denoted as re-divisions. These cells showed both S-phase markers but unlike DLS they were arranged as doublets, i.e. two daughter cells close to each other. The observed re-division frequency of NSCs is significantly higher than expected from random: While only 1.9±0.7% of NSCs are in S-phase at a given time point, 14.4±8.1% of those re-enter S-phase (p=7.8·10^−13^, two-sample Kolmogorov-Smirnov test, Figure 4U). We also observed re-divisions in *gfap*:GFP-cells, which were clearly distinguishable from each *gfap*:GFP+ cells (Supplementary Figure 4A) and occured in similar proportions (Supplementary Figure 4J). Notably, the *gfap:*GFP marker was the original label of the NSCs, without additional immunostaining for GFP, thus truly representing NSCs. Hence, dividing NSCs display a high likelihood of undergoing another division within the following days.

**Figure 4:**
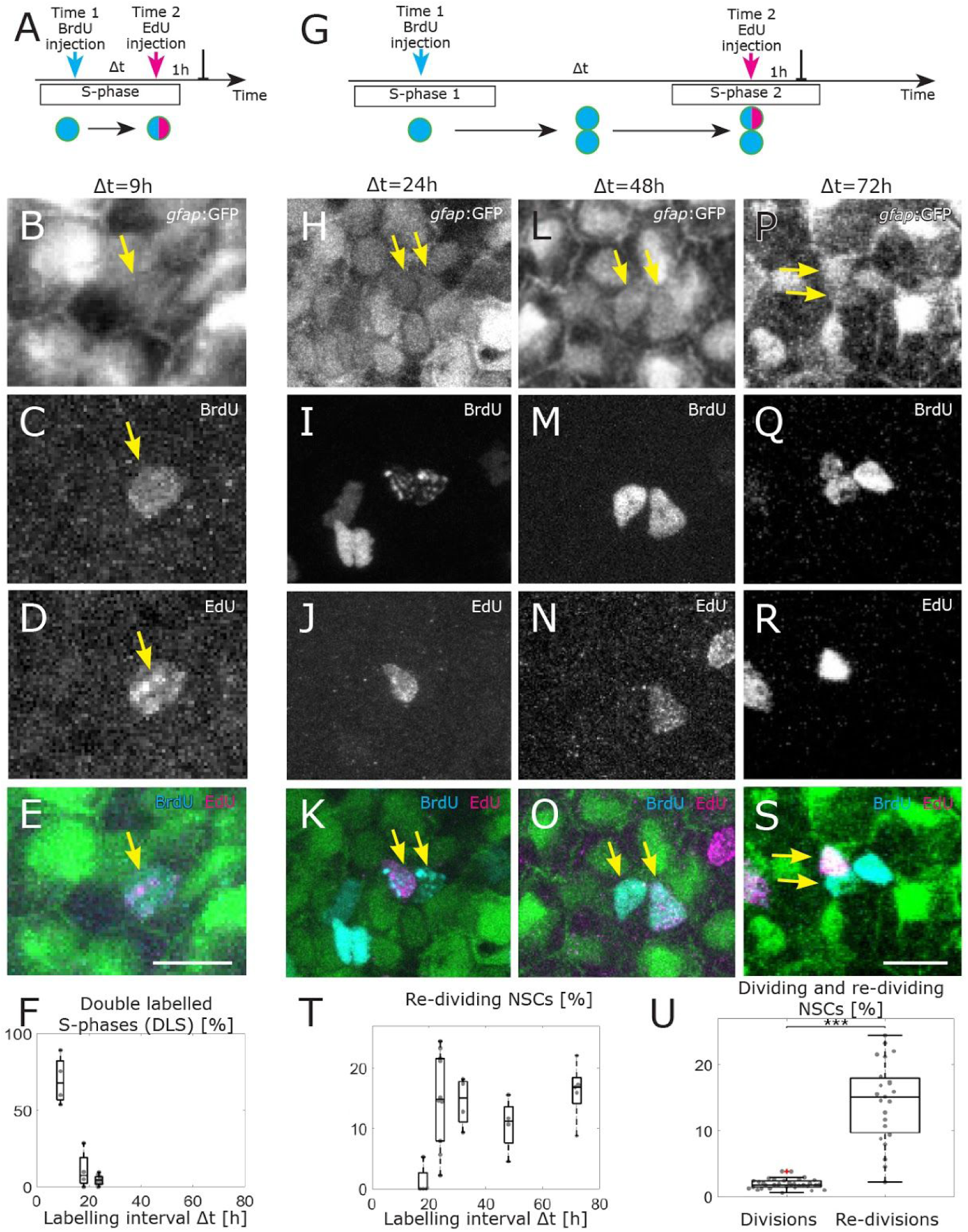
A large proportion of NSCs re-divide within 24 to 72 hours. (A) The two injections can label the same S-phase for small labelling intervals, leading to NSCs that are both EdU and BrdU positive, denoted as double-labelled S-phases (DLS). (B-E) Example DLS (yellow arrow) for a labelling interval Δt=9h. (F) The DLS proportion is high for a Δt of 9h and decreases rapidly with increasing Δt. Each dot represents the value for one brain hemisphere. (G) After a division, one of the daughter cells already labelled by the Time 1 label can enter in a new S-phase and incorporate a second label. This cell thereby re-divides. (H-S) Three examples of re-dividing NSCs with labelling intervals of 24h (H-K), 48h (L-O) and 72h (P-S). (T) The proportions of re-dividing NSCs within the dividing NSCs at Time 1 remain high from 24h to 72h labelling intervals. Each dot represents the value of one brain hemisphere. (U) The mean proportion of re-dividing NSCs (14.4 ± 8.1%) significantly (p-value=7.8·10^−13^, two-sample Kolmogorov-Smirnov test) exceeds the proportion of S-phase NSCs within all NSCs (1.9 ± 0.7%). Scale bar 10μm. Box plots range from the 25th to the 75th percentile, the central mark indicates the median and whiskers include points that are no outliers.

To confirm our observations, we performed independent experiments with a third time point of observation. We injected first BrdU, then EdU 24h later, and dissected the fish another 24h later, adding a PCNA immunostaining as a third cell cycle marker (Supplementary Figure 4B-I). In those brains, we found all combinations of NSC re-divisions: BrdU+ daughter cells that incorporated EdU, BrdU+ daughter cells that expressed PCNA, and EdU+ daughter cells that expressed PCNA. EdU+ doublets, which were also PCNA+ might represent cells that reached the end of a cell cycle without necessarily re-entering a second division round. However, we also observed EdU+ doublets in which only one daughter cell was PCNA+, signifying a specific cell cycle re-entry of this daughter cell.

We could not assess how many rounds of reiterated divisions NSCs may undergo maximally, as an increasing number of BrdU-labelled cells with an increasing chase time renders a discrimination between neighboring clones impossible. However, experiments carried out with 72 hours labelling interval revealed the presence of triplets of BrdU-labelled *gfap*:GFP+ cells (Figure 4Q), indicating that at least two rounds of divisions have been taking place within this time window. This result implies that the distribution of cell cycle entries in the NSC population is also a result of the recent history of individual cells.

### An agent based model of re-dividing NSCs recreates aggregated spatio-temporal patterns observed experimentally

To quantitatively evaluate if re-divisions suffice to induce the observed aggregated patterns, we simulated dividing NSCs with an agent based spatio-temporal model. However, fitting a spatio-temporal model to data is extremely challenging, since parameter estimation relies on repeated, computationally expensive simulations <u>(Jagiella et al., 2017)</u>. We thus split our simulation approach into two steps: First we fit a simple, non-spatial model to the observed DLS and re-dividing NSCs for different labelling intervals Δt (Figure 4F,T) to derive the length and variability of cell cycle and S-phase. Second, we simulate an agent based spatio-temporal model with the inferred parameters, analyze the simulated patterns statistically, and compare the results with experimental data.

To infer cell cycle kinetics from our data we implemented a non-spatial model of dividing NSCs with 5 parameters: the minimal cell cycle length d_cc_ and minimal S-phase length d_sp_, their variability *β*_cc_ and *β*_sp_ parametrizing a lag-exponential <u>(Weber et al., 2014)</u> distribution, and the re-division probability p_re-div_ (see Supplementary Figure 5A). From this model we simulate dividing NSCs and a BrdU-EdU-labelling experiment (Supplementary Figure 5B) and evaluate the percentage of re-dividing cells and DLS. We optimized the parameters of our model to the observed frequencies (see Methods) using approximate Bayesian computation <u>(Klinger et al., 2018)</u>. Our model fitted the data, in particular the plateau of re-dividing cells (Supplementary Figure 5C), and the sharp decrease of DLSs after 9h (Supplementary Figure 5D). It predicted a minimal cell cycle time of 22.2h with a mean cell cycle time of 107.5h, a minimal S-phase length of 16.6h with a mean S-phase length of 18.2, and a re-division probability of 0.38 (Supplementary Figure 5E,F).

**Figure 5:**
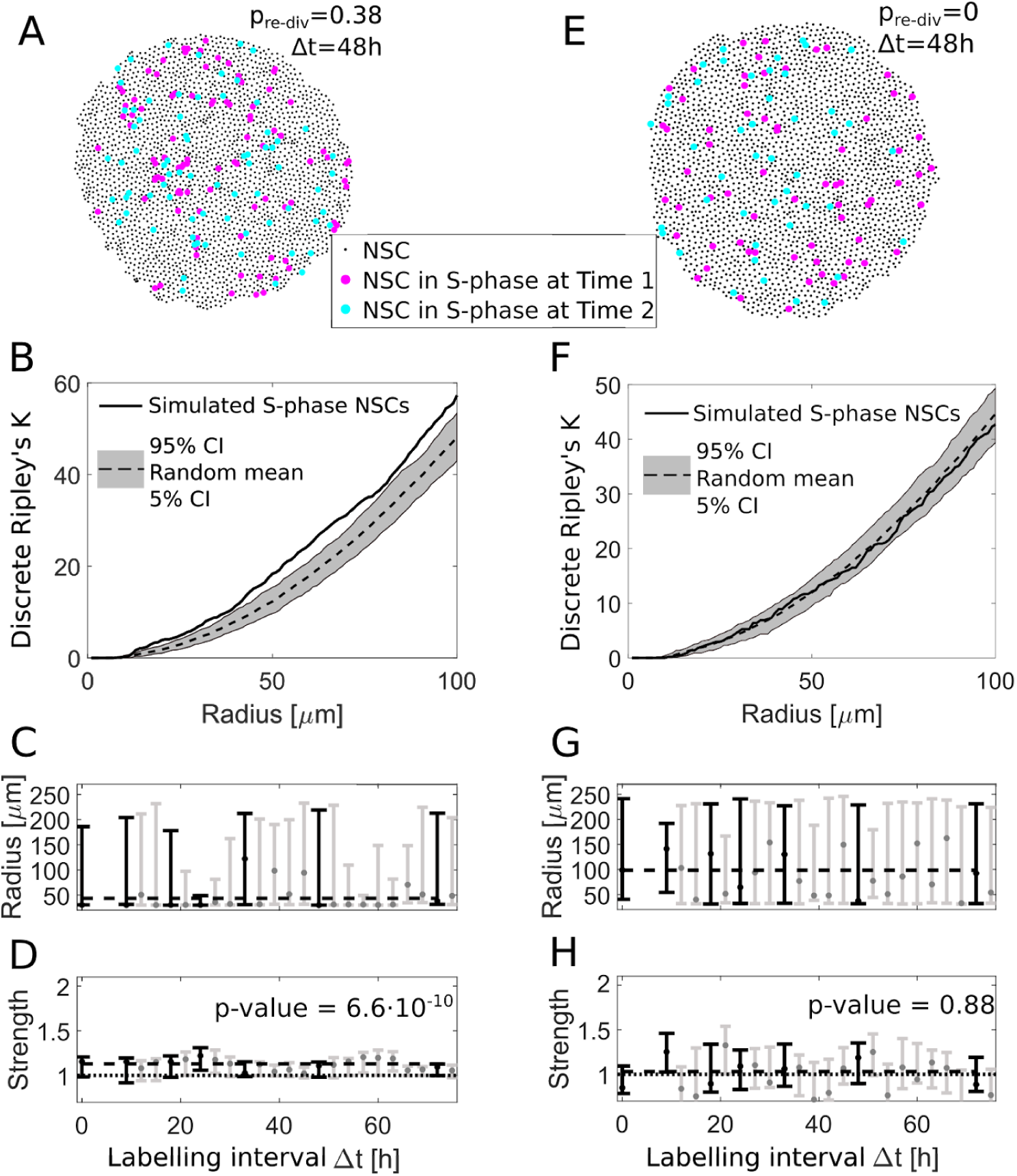
An agent based re-division model can explain spatial aggregation of S-phase NSCs. (A) We simulate NSC divisions with a re-division probability of p_re-div_=0.38 and take virtual measurements (here with Δt=48h). (B) The simulated S-phase NSCs in (A) exhibit an aggregated spatio-temporal pattern according to the discrete Ripley’s K curve (solid line) which is above the 90% confidence interval of randomly sampled patterns (grey area), similar to experimentally observed patterns. (C) The fitted radii for simulations with re-dividing NSCs for different labelling intervals Δt are variable with a maximum likelihood value of 50 μm. (D) The respective fitted strengths values are above 1 indicating aggregated patterns. Fitting a constant model to the most likely values with the same Δt as experimentally observed (black bars) reveals a significant shift (p-value = 6.6 ·10^−10^) from a strength of 1. (E) Simulated NSC divisions and virtual measurements with a re-division probability p_re-div_=0 at two labellings Δt=48h apart. (F) Without re-divisions, the simulated S-phase NSCs are within the boundaries of random patterns. (G) The fitted radii are highly variable with maximum likely values from 50 to 150 μm. (H) In contrast to the simulations with re-divisions, we find no indication for aggregated patterns for all labelling intervals (p-value = 0.88 for a constant model with non-zero shift from a strength of 1).

We now fed an agent based model, implemented in the Morpheus toolbox <u>(Starruß et al., 2014)</u> with the inferred parameters to generate spatio-temporal patterns (see Methods). After a transient phase, we simulated a first measurement by marking all cells in S-phase at that time point. We simulated NSC dynamics further for different labelling intervals and simulated a second measurement, analogously to the first. We then created a distance matrix for S-phase cells at the two timepoints (exemplary patterns shown in Figure 5A for Δt = 48h), calculated the discrete Ripley’s K (see Figure 5B) and inferred interaction radius and strength, analogously to experimental data processing. Analogously to our experimental data we found a significant (p-value= 6.6 10^−10^) positive interaction strength (Figure 5D), and an interaction radius around 45μm, which was a bit smaller than experimentally observed (Figure 5C).

In control simulations, we omitted re-divisions. In that case, all NSCs entered division with a probability of 2 10^−3^ per hour. In the corresponding simulated measurements (Figure 5E), neither the discrete Ripley’s K (see Figure 5F) nor interaction parameters hint to an aggregation of divisions. While the interaction radius strongly varied around a mean value of 89±42 μm (Figure 5G), the interaction strength is 0.97±0.2, indicating random division patterns (Figure 5H). Thus, with re-divisions and no additional spatial feedback, our agent based simulations created aggregated spatio-temporal patterns of dividing NSCs as observed experimentally.

## Discussion

We detected non-random, aggregated patterns of dividing NSCs in the zebrafish brain. This finding led us to hypothesize about causative spatio-temporal effects. The observation of re-dividing NSCs together with our agent based simulations confirmed that a group of cells, actively dividing for several days, is able to create the emerging aggregated division patterns.

The emergence of complex patterns from simple rules has been analyzed extensively, e.g. for artificial systems <u>(Marr and Hütt, 2005)</u>, biology inspired models <u>(Kauffman and Clayton, 2006)</u>, and biological phenomena <u>(Manukyan et al., 2017)</u>. Here, we abstract quiescent and dividing stem cells as agents in a continuous space, and use a stochastic model of the cell cycle dynamics to quantitatively compare two hypotheses: Do we need cell-cell interactions or niche effects to generate the observed aggreaged spatio-temporal division patterns, or do re-divisions of stem cells suffice? We find that the simplest model, i.e. one with no feedback but active, re-dividing NSCs is able to explain the emergent patterns observed in the zebrafish brains.

According to this model, coordination of cell cycle entries in the population would be regulated by the internal synchronization due to the individual cell history. Nevertheless we cannot reject the existence of additional mechanisms that might contribute to these patterns, such as local diffusive signaling activity in delimited groups of cells, a functional activity of extended cell-cell contacts, as has been observed on NSCs <u>(Obermann et al., 2019)</u>, or the presence of intermingled NSCs that are intrinsically less likely to enter cell cycle. The observed non-random spatial distribution indeed could suggest that some NSCs would be more susceptible to be recruited from the quiescent state than others. Some levels of heterogeneity have also been observed molecularly, for instance, variable levels of the Zinc finger protein Fezf2 regulate Notch activity levels and quiescence of NSCs <u>(Berberoglu et al., 2014)</u>. Likewise, the expression of miR9 involved in keeping quiescence upstream of Notch signaling is found only in a subset of quiescent cells <u>(Katz et al., 2016)</u> several days after a division. In a recent study, a subpopulation of NSCs that expresses low levels of Elavl3 has been characterized as mostly non-dividing cells in transit towards neuronal differentiation (Lange et al., 2020). Such differential expression levels within NSCs could hint towards a non-equipotent population. On the other hand, these differences might reflect fluctuating levels encountered equally in all cells. Seen that way, NSCs would be equipotent and therefore the non-random spatio-temporal distribution of S-phases that we observe emerges as the result of the re-divisions of daughter cells, creating specific spatial patterns. Hence, NSCs might firstly enter the cell cycle due to fluctuations of gene expression, as seen in culture <u>(Overton et al., 2014)</u>, but once activated would undergo several rounds of cell cycle before re-entering a quiescent mode.

Repeated divisions of NSCs were not expected, given the low percentage of divisions in the whole NSC population. NSCs were able to enter successive rounds of cell division within a few days, the earliest starting 24 hours after the previous S-phase. This contrasts with the early model of adult NSCs established in the mouse dentate gyrus and subependymal zone, where radial astrocytes are quiescent and can replenish the transient amplifying progenitor population after all fast dividing cells have been eradicated by an ARA-C treatment (Doetsch et al., 1999). The latest characterizations by single cell sequencing differentiate between quiescent versus active radial astrocytes, however these most probably represent alternating states, as also suggested by a continuum of transcriptional states (Llorens Bobadilla, 2015, Dulken et al., 2017). The definition of an active radial astrocytes here cannot distinguish between a repetitive or sporadic division behavior. Recently however, evidence for several rounds of divisions within a few days in NSCs have been reported in the mouse subependymal zone from clonal analysis data using the Troy-driven recombination <u>(Basak et al., 2018)</u>. Further, live imaging of recombined Ascl-driven recombination in the dentate gyrus in vivo demonstrated the existence of several divisions of radial astrocytes in a short time window <u>(Pilz et al., 2018)</u>. In zebrafish, two studies based on in vivo imaging of the whole brain have followed NSCs, one observing single-labelled NSCs and their behavior of division and differentiation <u>(Barbosa et al., 2015)</u>, the other considering the whole dividing NSC population with the help of two transgenic lines, her4:RFP and mcm5: GFP, highlighting the NSCs and the cell cycle, respectively <u>(Dray et al., 2015)</u> over the course of two weeks. Rapidly re-dividing daughter stem cells, as observed by our double labelling approach in living fish combined with confocal imaging that allows for a clear distinction between nuclei close to each other, have not been reported yet. It is likely that they have been missed due to the difficulty to resolve distinct sister cells with intravital imaging in previous studies.

How these recurrent divisions of adult stem cells come about will be important to assess in the future. Studies on human epithelial cell lines have shown that mother cells can relay distinct levels of CCND1 mRNA and p53 protein to their daughters, which after completing mitosis will then rapidly decide for the next round of division, depending on a resulting bimodal level of activity of cyclin dependent kinase (CDK2) <u>(Spencer et al., 2013; Yang et al., 2017)</u>. It could well be that in the NSCs too, specific cell cycle regulators would be transmitted to daughter cells, thereby permitting for immediate new rounds of division. Other mechanisms of daughter cells’ decisions for quiescence or new cycle have been reported, for instance, in the adult mouse dentate gyrus, degradation of Ascl1a in daughter cells mediated by the ubiquitin ligase Huwe1 takes place and promotes a re-entry in quiescence <u>(Urbán aet al., 2016)</u>.

Hence, in stem cell populations, individual phases of quiescence and exit thereof might well be predictable according to inherited factors, allowing us to understand how the dynamics tissue maintenance are regulated and how strong the level of stochasticity is.

## Data availability

We provide access to raw image data,processed single cell information, code for analysis, parameter inference and result figures at https://github.com/QSCD/spatiotemporalAnalyses.

## Methods

### Zebrafish maintenance and transgenic lines

Zebrafish of the transgenic line *gfap*:GFP (mi2001) <u>(Bernardos and Raymond, 2006)</u> were bred and maintained in the fish facility of the Helmholtz Zentrum München. Experiments were conducted under the animal protocol 55.2-1-2532-83-14, in accordance with animal welfare rules of the government of Oberbayern.

### Labelling of consecutive S-phases *in vivo*

5-bromo-2’-deoxyuridine (BrdU) or 5-Ethynyl-2’-deoxyuridine (EdU) were dissolved at a concentration of 1mg/ml in saline solution (0.07% NaCl) containing methylene blue and injected intraperitoneally (5ul/ 0.1g body mass) into the fish at precise time points. Fish were over-anasthetized in MS-222 or placed in ice-water for euthanasia, decapitated and the brains dissected and fixed in 4% PFA overnight. Whole mount brains were processed for Click chemistry with Azyde-Alexa-Fluor-647 to detect EdU, following the manufacturer’s protocol (Thermofischer C-10269), see also <u>(Qu et al., 2011)</u>. For the subsequent BrdU immunoreaction, brains were treated with HCl 2M at 37°C for 30 minutes, washed in sodium tetraborate buffer 0.1M, pH8 and in PBS, and incubated with Mouse-anti BrdU (Phoenix PRB1-U) 1:800 overnight. Mouse or Rabbit anti PCNA (clone PC-10, Santa Cruz sc-56; Abcam ab15497) was diluted 1:800 or 1:100 respectively. Following secondary antibodies were incubated for 1h at room temperature at 1:1000 concentration: Goat-anti-mouse Alexa-Fluor-555, Goat-anti-mouse Alexa-Fluor-405, Goat-anti-Rabbit-Pacific Blue, Goat-anti-Rabbit Alexa-Fluor-555 (Thermofischer).

### Imaging of whole mount brains

Brains were mounted in vectashield or in PBS with 40% glycerol between two coverslips separated by 8-layered-parafilm spacers. Imaging was performed using a Leica SP5 confocal microscope with a 20x glycerol immersion objective at a resolution of 2048 × 2048 pixels, and for close up views with a 63x glycerol immersion objective.

### Identification of *gfap*:GFP+ cells and PCNA/EdU/BrdU-labelled cells

NSCs that are labelled by GFP in the transgenic *gfap*:GFP line were identified using the Single Cell Identification Pipeline (SCIP, <u>(Lupperger et al., 2018)</u>). In this pipeline single cells are automatically identified from an image 3D stack, exploiting the fact that all NSCs are located on top of the hemisphere on a 2D surface. SCIP returns x, y and z coordinates for all NSCs. Nuclei labelled by PCNA immunochemistry, BrdU immunochemistry or by EdU-click chemistry were identified semi-automatically: nuclei were first identified with SCIP and then visually verified one by one to avoid false positives and ensure correct assignment to GFP-positive or negative cells, using the two channels in consecutive z-planes of the confocal stacks.

In experiments with Δt>18h, the majority of Time 1-labelled cells were found as doublets, i.e. a pair of small cells close to each other (see Figure 4K,O,S), since the timespan allowed the mother cell to reach its mitotic (M) phase. For Δt=72h even triplets occured. Doublets and triplets were handled as single division events in the subsequent analysis. Some of the doublets incorporated the second dU label and were categorized only to Time 1 in the spatio-temporal analysis, in order to assess distances specifically between the distinct sets of cells undergoing S-phases at consecutive time points. The proportion of re-dividing NSCs was calculated by dividing the number of doublets (or triplets) containing at least one GFP+BrdU+EdU+ cell by the total number of GFP+BrdU+ doublets (or triplets).

### Spatial statistics

Spatial statistics were applied on NSCs labelled in S-phase, i.e. *gfap:*GFP+ cells with EdU or BrdU staining. The set of NSCs without EdU or BrdU staining served as the substrate to e.g. evaluate patterns for randomly dividing NSCs.

#### 3D Distance matrix

We determine the distance between any two cells by calculating the shortest path on the hemisphere manifold to account for the bending of the hemisphere surface. To this end we used the fitted hemisphere surface and calculated the stepwise shortest paths between two cell locations on it (see <u>(Lupperger et al., 2018)</u> for a detailed description).

#### Ripley’s K statistics

Ripley’s K <u>(Ripley, 1976)</u> is a measure for the deviation of a point pattern from spatial homogeneity. It has been previously applied in geographic information science <u>(Hohl et al., 2017)</u>, in the context of spatial economic analysis <u>(Arbia et al., 2017)</u> and archaeological studies <u>(Negre et al., 2018)</u>. In all those applications, it is assumed that the point pattern occurs from a Poisson point process on a homogenous space. Here, however, division events can only occur where NSCs are located on the 2D hemisphere manifold. We adapted the measure to account for this inhomogeneity by sampling a point pattern only from discrete NSC locations and thus call it discrete Ripley’s K.

For *S* S-phase NSCs we calculate the spatial *K*_*S*_ value for increasing radii *r* along

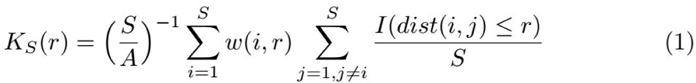

This function counts NSCs in S-phase within a radius *r* around the S-phase NSC *i*. The indicator function *I* is 1 if cell *i* and *j* are closer than *r* and 0 otherwise. The term is normalized by the total amount of NSCs in S-phase *S* and by the S-phase NSC density *S*/*A*. The hemisphere area *A* is calculated as the sum of all triangle areas between all NSCs obtained via delaunay triangulation (see Supplement Figure 1A) <u>(Delaunay and Others, 1934)</u>. The edge correction term *w*(*i,r*) is 1 if the disc with radius *r* around NSC *i* does not cut the hemisphere edge and else calculated as the fraction of disc inside the hemisphere.

To obtain the spatio-temporal *K*_*ST*_ between two sets of NSCs in S-phase, labelled at Time 1 and Time 2 we modified equation 1:

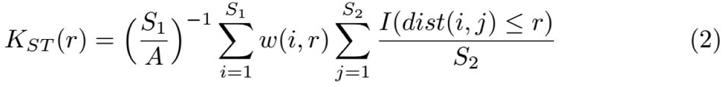

We here count *S*_2_ (NSCs in S-phase at Time 2) cells around *S*_1_ (NSCs in S-phase at Time 1) cells within r. In contrast to equation 1 the whole term is normalized by the density of *S*_1_ cells while the indicator function is divided by the number of *S*_2_ cells.

To compare observed K values to random spatial distributions we sample the amount of observed S and *S*_2_ cells, respectively, (*S*_1_ cell locations are fixed) from all NSCs 20 times and calculate the random sampling discrete K value. To evaluate whether the observed K value differs from random sampling we check if the observed K value is below the 5% quantile (which would indicate spatial dispersion) or above the 95% quantile (which would indicate spatial aggregation) of the 20 sampled discrete K values.

To make Ripley’s K plots comparable between hemispheres, we standardized the results per hemisphere similar to a z-score, such that K values are 1 when they are on the 95% quantile and -1 when they are on the 5% quantile.

### Model based analysis

#### Influence model

To determine the spatial extent and the nature of the temporal influence of divisions at different timepoints, we extend a spatial influence model <u>(Lupperger et al., 2018)</u> with two parameters: the influence strength (*g*), where *g*=1 means no dependencies, below 1 dispersion between the two populations, and above 1 aggregation. The second parameter the model fits is the influence radius (*r*) in μm.

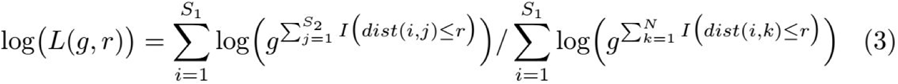

For each hemisphere, we fit the parameters (*g,r*) of the model to locations of cells in S-phase of two time points to detect spatial dependencies of NSCs in S-phase at the later time point (*S*_2_) regarding NSCs in S-phase at the earlier time point (*S*_1_). The parameters are fitted with a log-likelihood approach (Eq. 3) where *S*_x_ is the number of cells in S-phase at time point x, while *N* is the whole NSC population. The indicator function *I* is 1 if the distance between two cells (*i, j* or *i, k*) is smaller or equal *r*.

In the numerator term we iterate over all *S*_1_ cells and count the number of *S*_2_ cells having a smaller or equal distance than r, while in the denominator we normalize the equation by counting the not affected (non-S-phase) cells of N within r around *S*_1_ cells.

Equation 3 calculates the log likelihood for one hemisphere. To calculate the log likelihood across several hemispheres every single likelihood per hemisphere is summed up and optimized in parallel to form a combined likelihood.

We use the PESTO toolbox <u>(Stapor et al., 2018)</u> to maximize the likelihood including uncertainty estimation via posterior sampling <u>(Ballnus et al., 2018)</u> as can be seen exemplarily in Figure 3B for hemispheres with Δt=48h and for all hemispheres in Supplementary Figure 3.

#### Cell division model

The observed fractions of re-dividing NSCs and DLS (Fig. 4S,T) suggest an upper limit for the S-phase length of 32h (since we observe no DLSs for Δt ≥ 32h, Fig. 4S) and a lower limit for all other cell cycle phases of 9h (since we observe no re-diving cells at Δt = 9h, Fig. 4T). A simple model with a fixed cell cycle length of e.g. 32h+9h is however not able to explain a plateauing of reviding NSCs for Δt ≥ 24h. Instead of constant cell cycle and S-phase length, we thus assume them to be distributed as delayed exponential distributions <u>(Weber et al., 2014)</u>,

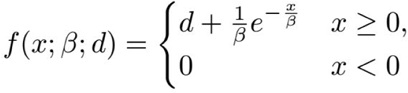

where *β* is the scale parameter of the exponential distribution and *d* is the delay (see Supplementary Figure 5A). Additionally we assume a re-division probability *p*_re-div_, which is lower bounded by the observed re-division frequency of 15%, since we are not able to observe all re-divisions (compare Supplementary Figure 4J and Supplementary Figure 5B) in our snapshot experiments with an EdU labelling of 1 hour.

To infer these five parameters (scale parameter and delay of cell cycle, *d*_cc_ and *β*_cc_, and S-phase, *d*_sp_ and *β*_sp_, and re-division probability *p*_*re-div*_) from (i) the observed re-division frequency and (ii) the fraction of DLS NSCs (see sketches in Supplementary Figure 5C,D) at every labelling interval Δt=9h, 18h, 24h, 32h, 48h, and 72h, we apply approximate Bayesian computation (ABC) <u>(Klinger et al., 2018)</u>. Given the five parameters (*d*_*cc*_ *β*_*cc*_ *d*_*sp*_ *β*_*sp*_ and *p*_*re-div*_) we simulate the dynamics of 10000 dividing cells over 400-500h. Using a fixed endpoint (Time 2) we can determine Time 1 by subtracting the according labelling interval. With two virtual measurements we calculate the observed re-division and double labelled S-phase (DLS) frequency via counting all cells being in S-phase at Time 1 and determine the status of these Time 1 S-phase cells at Time 2 (see Supplementary Figure 5B): If an S-phase of a cell is labelled by both virtual measurements, it is classified as DLS and if an offspring cell is is labelled in S-phase the cell is classified as re-dividing cell. The simulated re-division frequency is then the number of re-dividing cells divided by number of S-phase cells at Time 1 and the DLS fraction is number of DLSs divided by number of S-phase cells at Time 1, accordingly.

In order to fit the observed data we define a distance function between the observed and simulated re-division and DLS fractions per labelling interval. To this end, we calculate the sum of differences between means and standard deviations respectively:

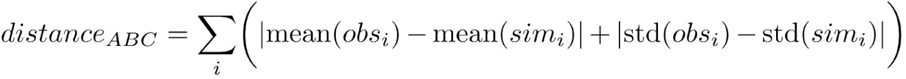

where i ∈ {9h, 18h, 24h, 32h, 48h, and 72h}. We optimize this distance function employing ABC with 50 epochs, evaluating 500 particles per epoch (ABC parameters).

### Agent based spatio-temporal simulation

To assess the contribution of re-division events to the emerging spatial patterns of NSCs in S-phase, we simulated an NSC population using the Morpheus algorithm <u>(Starruß et al., 2014)</u>. Morpheus is a modeling and simulation environment where cells act and interact as agents in space and time. We fix the base division probability of all cells to p_div_ = 9•10^−4^ per hour. With a mean S-phase length of 18h (inferred from cell division model above), this leads to a dividing cell fraction of 9•10^−4^ 1/h x 18h = 1.6% at every snapshot. NSCs re-dividing with a probability of p_re-div_ = 0.38 (inferred from cell division model above) increase this fraction (as they divide after one cell cycle independently of the base division probability) to roughly 2%, in line with experimental data (observed “Divisions” in Figure 4U).

We estimate the rate of NSC differentiation from the proportion of doublets with one *gfap*:GFP+ and one *gfap*:GFP-cell. This proportion is roughly 10% in our data. *Gfap*:GFP-cells are simulated for one more cell cycle duration and are then excluded, mimicking differentiated cells transitioning away from the stem cell pool inside the brain (Rothenaigner et al., 2011).

Cell cycle length is inferred from the cell division model (see above) with delay *d*_*cc*_ and *β*_*cc*_ from the delayed exponential distribution. Other predefined parameters are minimum and maximum cell size, determined by measuring real NSC size and simulated via sigmoidal cell growth. Morpheus implements basic cell-cell interactions and kinetic assumptions <u>(Starruß et al., 2014)</u>.

Our simulation starts with 500 cells and runs until the colony size reaches the observed 2356±460 (mean±sd) cells (see Figure 5A,E). We analyze cells in S-phase at the simulated measurement time points and apply the same spatial statistics as for the real data.

## Acknowledgements

We thank Carolin Loos and Lisa Bast for their advice regarding parameter inference, Yannik Schälte for his support with ABC, Benjamin Ballnus for his insights in MCMC sampling, Elmar Spiegel and Hannah Busen for statistical consulting and the rest of the Marr lab for helpful comments. Also we thank Walter de Back for his help to set up morpheus and Jovica Ninkovic for his valuable remarks. Furthermore we thank Moritz Thomas for their notes on the mascript and Elly Pachel for her help on code/analysis reproducibility. V.L. was funded by the BMBF, grant 01ZX1710A-F (Micmode-I2T). The experimental work was carried out in the lab of Hernán Lopez-Schier.

## Author contributions

V.L. processed the data and performed analyses. P.C. conducted all experiments. V.L., C.M. and P.C. wrote the article. C.M. and P.C. conceived and supervised the study.

## References

Alunni, A., Krecsmarik, M., Bosco, A., Galant, S., Pan, L., Moens, C.B., and Bally-Cuif, L. (2013). Notch3 signaling gates cell cycle entry and limits neural stem cell amplification in the adult pallium. Development 140, 3335–3347.

Arbia, G., Espa, G., Giuliani, D., and Dickson, M.M. (2017). Effects of missing data and locational errors on spatial concentration measures based on Ripley’s K-function. Spatial Economic Analysis 12, 326–346.

Ballnus, B., Schaper, S., Theis, F.J., and Hasenauer, J. (2018). Bayesian parameter estimation for biochemical reaction networks using region-based adaptive parallel tempering. Bioinformatics 34, i494–i501.

Barbosa, J.S., Sanchez-Gonzalez, R., Di Giaimo, R., Baumgart, E.V., Theis, F.J., Götz, M., and Ninkovic, J. (2015). Neurodevelopment. Live imaging of adult neural stem cell behavior in the intact and injured zebrafish brain. Science 348, 789–793.

Basak, O., Krieger, T.G., Muraro, M.J., Wiebrands, K., Stange, D.E., Frias-Aldeguer, J., Rivron, N.C., van de Wetering, M., van Es, J.H., van Oudenaarden, A., et al. (2018). Troy+ brain stem cells cycle through quiescence and regulate their number by sensing niche occupancy. Proc. Natl. Acad. Sci. U. S. A. 115, E610–E619.

Berberoglu, M.A., Dong, Z., Li, G., Zheng, J., Trejo Martinez, L. del C.G., Peng, J., Wagle, M., Reichholf, B., Petritsch, C., Li, H., et al. (2014). Heterogeneously expressed fezf2 patterns gradient Notch activity in balancing the quiescence, proliferation, and differentiation of adult neural stem cells. J. Neurosci. 34, 13911–13923.

Bernardos, R.L., and Raymond, P.A. (2006). GFAP transgenic zebrafish. Gene Expression Patterns 6, 1007–1013.

Chapouton, P., Skupien, P., Hesl, B., Coolen, M., Moore, J.C., Madelaine, R., Kremmer, E., Faus-Kessler, T., Blader, P., Lawson, N.D., et al. (2010). Notch activity levels control the balance between quiescence and recruitment of adult neural stem cells. J. Neurosci. 30, 7961–7974.

Chapouton, P., Webb, K.J., Stigloher, C., Alunni, A., Adolf, B., Hesl, B., Topp, S., Kremmer, E., and Bally-Cuif, L. (2011). Expression of hairy/enhancer of split genes in neural progenitors and neurogenesis domains of the adult zebrafish brain. J. Comp. Neurol. 519, 1748–1769.

Delaunay, B., and Others (1934). Sur la sphere vide. Izv. Akad. Nauk SSSR, Otdelenie Matematicheskii I Estestvennyka Nauk 7, 1–2.

Dong, J., Pan, Y.-B., Wu, X.-R., He, L.-N., Liu, X.-D., Feng, D.-F., Xu, T.-L., Sun, S., and Xu, N.-J. (2019). A neuronal molecular switch through cell-cell contact that regulates quiescent neural stem cells. Sci Adv 5, eaav4416.

Dray, N., Bedu, S., Vuillemin, N., Alunni, A., Coolen, M., Krecsmarik, M., Supatto, W., Beaurepaire, E., and Bally-Cuif, L. (2015). Large-scale live imaging of adult neural stem cells in their endogenous niche. Development 142, 3592–3600.

Edelmann, K., Glashauser, L., Sprungala, S., Hesl, B., Fritschle, M., Ninkovic, J., Godinho, L., and Chapouton, P. (2013). Increased radial glia quiescence, decreased reactivation upon injury and unaltered neuroblast behavior underlie decreased neurogenesis in the aging zebrafish telencephalon. J. Comp. Neurol. 521, 3099–3115.

Ehm, O., Göritz, C., Covic, M., Schäffner, I., Schwarz, T.J., Karaca, E., Kempkes, B., Kremmer, E., Pfrieger, F.W., Espinosa, L., et al. (2010). RBPJkappa-dependent signaling is essential for long-term maintenance of neural stem cells in the adult hippocampus. J. Neurosci. 30, 13794–13807.

Folgueira, M., Bayley, P., Navratilova, P., Becker, T.S., Wilson, S.W., and Clarke, J.D.W. (2012). Morphogenesis underlying the development of the everted teleost telencephalon. Neural Dev. 7, 32.

Furlan, G., Cuccioli, V., Vuillemin, N., Dirian, L., Muntasell, A.J., Coolen, M., Dray, N., Bedu, S., Houart, C., Beaurepaire, E., et al. (2017). Life-Long Neurogenic Activity of Individual Neural Stem Cells and Continuous Growth Establish an Outside-In Architecture in the Teleost Pallium. Curr. Biol. 27, 3288–3301.e3.

Hohl, A., Zheng, M., Tang, W., Delmelle, E., and Casas, I. (2017). Spatiotemporal Point Pattern Analysis Using Ripley’s K Function. Geospatial Data Science Techniques and Applications 155–176.

Jagiella, N., Rickert, D., Theis, F.J., and Hasenauer, J. (2017). Parallelization and High-Performance Computing Enables Automated Statistical Inference of Multi-scale Models. Cell Systems 4, 194–206.e9.

Katz, S., Cussigh, D., Urbán, N., Blomfield, I., Guillemot, F., Bally-Cuif, L., and Coolen, M. (2016). A Nuclear Role for miR-9 and Argonaute Proteins in Balancing Quiescent and Activated Neural Stem Cell States. Cell Rep. 17, 1383–1398.

Kauffman, S., and Clayton, P. (2006). On emergence, agency, and organization. Biology and Philosophy 21, 501–521.

Klinger, E., Rickert, D., and Hasenauer, J. (2018). pyABC: distributed, likelihood-free inference. Bioinformatics 34, 3591–3593.

Knobloch, M., Braun, S.M.G., Zurkirchen, L., von Schoultz, C., Zamboni, N., Araúzo-Bravo, M.J., Kovacs, W.J., Karalay, O., Suter, U., Machado, R.A.C., et al. (2013). Metabolic control of adult neural stem cell activity by Fasn-dependent lipogenesis. Nature 493, 226–230.

Knobloch, M., Pilz, G.-A., Ghesquière, B., Kovacs, W.J., Wegleiter, T., Moore, D.L., Hruzova, M., Zamboni, N., Carmeliet, P., and Jessberger, S. (2017). A Fatty Acid Oxidation-Dependent Metabolic Shift Regulates Adult Neural Stem Cell Activity. Cell Rep. 20, 2144–2155.

Kroehne, V., Freudenreich, D., Hans, S., Kaslin, J., and Brand, M. (2011). Regeneration of the adult zebrafish brain from neurogenic radial glia-type progenitors. Development 138, 4831–4841.

Lepko, T., Pusch, M., Müller, T., Schulte, D., Ehses, J., Kiebler, M., Hasler, J., Huttner, H.B., Vandenbroucke, R.E., Vandendriessche, C., et al. (2019). Choroid plexus-derived miR-204 regulates the number of quiescent neural stem cells in the adult brain. EMBO J. 38, e100481.

Liang, J., Balachandra, S., Ngo, S., and O’Brien, L.E. (2017). Feedback regulation of steady-state epithelial turnover and organ size. Nature 548, 588–591.

Lupperger, V., Buggenthin, F., Chapouton, P., and Marr, C. (2018). Image analysis of neural stem cell division patterns in the zebrafish brain. Cytometry A 93, 314–322.

Manukyan, L., Montandon, S.A., Fofonjka, A., Smirnov, S., and Milinkovitch, M.C. (2017). A living mesoscopic cellular automaton made of skin scales. Nature 544, 173–179.

Marr, C., and Hütt, M.T. (2005). Topology regulates pattern formation capacity of binary cellular automata on graphs. Physica A 354, 641–662.

März, M., Chapouton, P., Diotel, N., Vaillant, C., Hesl, B., Takamiya, M., Lam, C.S., Kah, O., Bally-Cuif, L., and Strähle, U. (2010). Heterogeneity in progenitor cell subtypes in the ventricular zone of the zebrafish adult telencephalon. Glia 58, 870–888.

Mesa, K.R., Kawaguchi, K., Cockburn, K., Gonzalez, D., Boucher, J., Xin, T., Klein, A.M., and Greco, V. (2018). Homeostatic Epidermal Stem Cell Self-Renewal Is Driven by Local Differentiation. Cell Stem Cell 23, 677–686.e4.

Morrison, S.J., and Scadden, D.T. (2014). The bone marrow niche for haematopoietic stem cells. Nature 505, 327–334.

Mueller, T., and Wullimann, M.F. (2009). An evolutionary interpretation of teleostean forebrain anatomy. Brain Behav. Evol. 74, 30–42.

Negre, J., Muñoz, F., and Barceló, J.A. (2018). A Cost-Based Ripley’s K Function to Assess Social Strategies in Settlement Patterning. Journal of Archaeological Method and Theory 25, 777–794.

Obermann, J., Wagner, F., Kociaj, A., Zambusi, A., Ninkovic, J., Hauck, S.M., and Chapouton, P. (2019). The Surface Proteome of Adult Neural Stem Cells in Zebrafish Unveils Long-Range Cell-Cell Connections and Age-Related Changes in Responsiveness to IGF. Stem Cell Reports 12, 258–273.

Ottone, C., Krusche, B., Whitby, A., Clements, M., Quadrato, G., Pitulescu, M.E., Adams, R.H., and Parrinello, S. (2014). Direct cell–cell contact with the vascular niche maintains quiescent neural stem cells. Nature Cell Biology 16, 1045–1056.

Overton, K.W., Spencer, S.L., Noderer, W.L., Meyer, T., and Wang, C.L. (2014). Basal p21 controls population heterogeneity in cycling and quiescent cell cycle states. Proc. Natl. Acad. Sci. U. S. A. 111, E4386–E4393.

Pellegrini, E., Mouriec, K., Anglade, I., Menuet, A., Le Page, Y., Gueguen, M.-M., Marmignon, M.-H., Brion, F., Pakdel, F., and Kah, O. (2007). Identification of aromatase-positive radial glial cells as progenitor cells in the ventricular layer of the forebrain in zebrafish. J. Comp. Neurol. 501, 150–167.

Pilz, G.-A., Bottes, S., Betizeau, M., Jörg, D.J., Carta, S., Simons, B.D., Helmchen, F., and Jessberger, S. (2018). Live imaging of neurogenesis in the adult mouse hippocampus. Science 359, 658–662.

Qu, D., Wang, G., Wang, Z., Zhou, L., Chi, W., Cong, S., Ren, X., Liang, P., and Zhang, B. (2011). 5-Ethynyl-2’-deoxycytidine as a new agent for DNA labeling: detection of proliferating cells. Anal. Biochem. 417, 112–121.

Ripley, B.D. (1976). The second-order analysis of stationary point processes. J. Appl. Probab. 13, 255–266.

Rothenaigner, I., Krecsmarik, M., Hayes, J.A., Bahn, B., Lepier, A., Fortin, G., Götz, M., Jagasia, R., and Bally-Cuif, L. (2011). Clonal analysis by distinct viral vectors identifies bona fide neural stem cells in the adult zebrafish telencephalon and characterizes their division properties and fate. Development 138, 1459–1469.

Spencer, S.L., Cappell, S.D., Tsai, F.-C., Overton, K.W., Wang, C.L., and Meyer, T. (2013). The proliferation-quiescence decision is controlled by a bifurcation in CDK2 activity at mitotic exit. Cell 155, 369–383.

Stapor, P., Weindl, D., Ballnus, B., Hug, S., Loos, C., Fiedler, A., Krause, S., Hroß, S., Fröhlich, F., Hasenauer, J., et al. (2018). PESTO: Parameter EStimation TOolbox. Bioinformatics 34, 705–707.

Starruß, J., de Back, W., Brusch, L., and Deutsch, A. (2014). Morpheus: a user-friendly modeling environment for multiscale and multicellular systems biology. Bioinformatics 30, 1331–1332.

Than-Trong, E., and Bally-Cuif, L. (2015). Radial glia and neural progenitors in the adult zebrafish central nervous system. Glia 63, 1406–1428.

Urbán, N., van den Berg, D.L.C., Forget, A., Andersen, J., Demmers, J.A.A., Hunt, C., Ayrault, O., and Guillemot, F. (2016). Return to quiescence of mouse neural stem cells by degradation of a proactivation protein. Science 353, 292–295.

van Velthoven, C.T.J., and Rando, T.A. (2019). Stem Cell Quiescence: Dynamism, Restraint, and Cellular Idling. Cell Stem Cell 24, 213–225.

Weber, T.S., Jaehnert, I., Schichor, C., Or-Guil, M., and Carneiro, J. (2014). Quantifying the length and variance of the eukaryotic cell cycle phases by a stochastic model and dual nucleoside pulse labelling. PLoS Comput. Biol. 10, e1003616.

Yang, H.W., Chung, M., Kudo, T., and Meyer, T. (2017). Competing memories of mitogen and p53 signalling control cell-cycle entry. Nature 549, 404–408.

Yeh, C.-Y., Asrican, B., Moss, J., Quintanilla, L.J., He, T., Mao, X., Cassé, F., Gebara, E., Bao, H., Lu, W., et al. (2018). Mossy Cells Control Adult Neural Stem Cell Quiescence and Maintenance through a Dynamic Balance between Direct and Indirect Pathways. Neuron 99, 493–510.e4.

