## Supplemental information for "Aggregated spatio-temporal division patterns emerge from reoccurring divisions of neural stem cells"

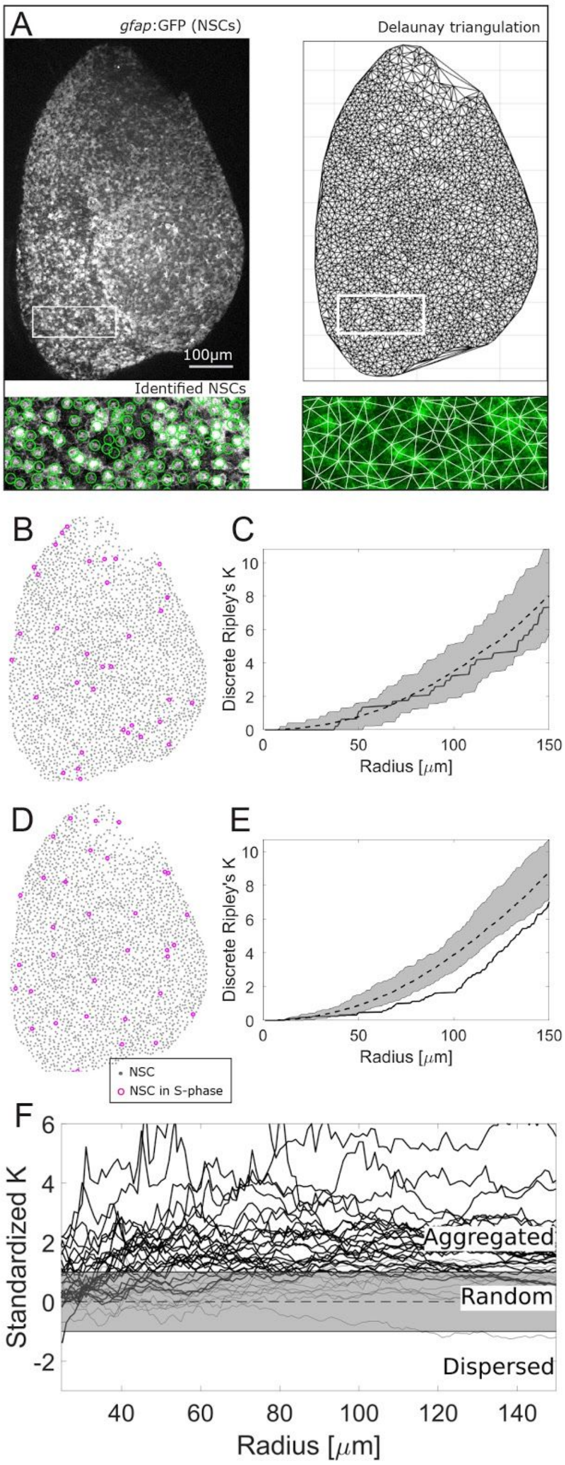


**Supplementary Figure 1.**

(A) Delaunay triangulation applied on the NSCs to calculate discrete Ripley´s K.
(B) A synthetically generated random pattern.
(C) Discrete Ripley’s K correctly identifies that pattern to be within the 90% confidence interval of randomly sampled patterns.
(D) A synthetically generated dispersed pattern with radius 100 µm.
(E) Discrete Ripley’s K correctly identifies that pattern below the random range.
(F) Standardized K shows S-phase NSC aggregation in 25 (thick lines) out of 36 hemispheres. Hemispheres which show a mean standardized K value higher than 1 between 30 and 150µm are labeled with a thick line. Each trace represents the spatial distribution of S-phase NSCs in one hemisphere.

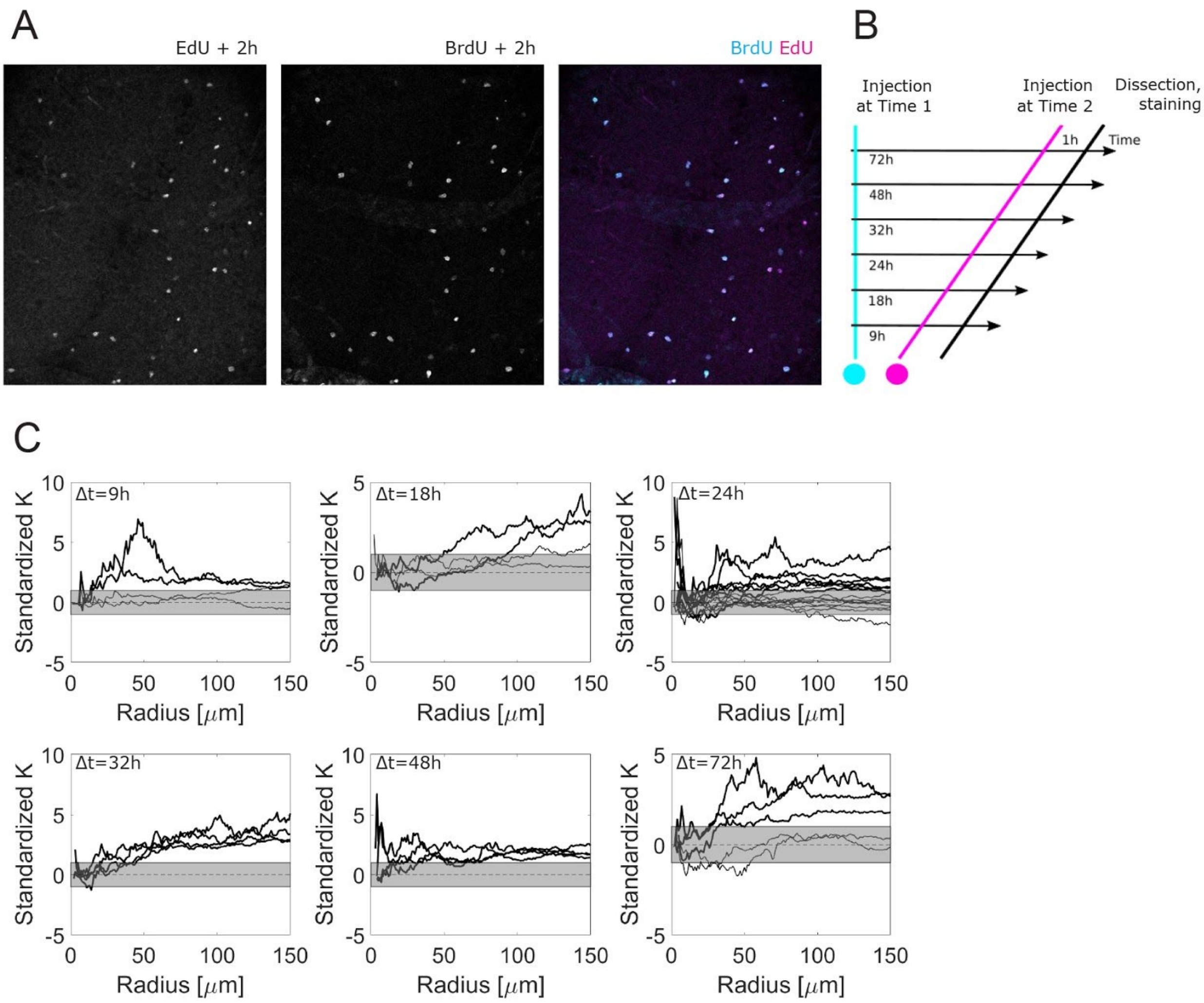


**Supplementary Figure 2.**

(A) Control EdU and BrdU labelling. EdU and BrdU have been injected simultaneously 2 hours before sacrificing the fish and therefore label the same set of cells.

(B) Experimental setup for measuring spatio-temporal patterns. With different labelling intervals $\Delta$t between the two S-phase-labellings we systematically profile spatio-temporal effects.

(C) Standardized Ripley’s K shows spatio-temporal aggregation of S-phase NSCs in 22 out of 36 hemispheres for different $\Delta$t.

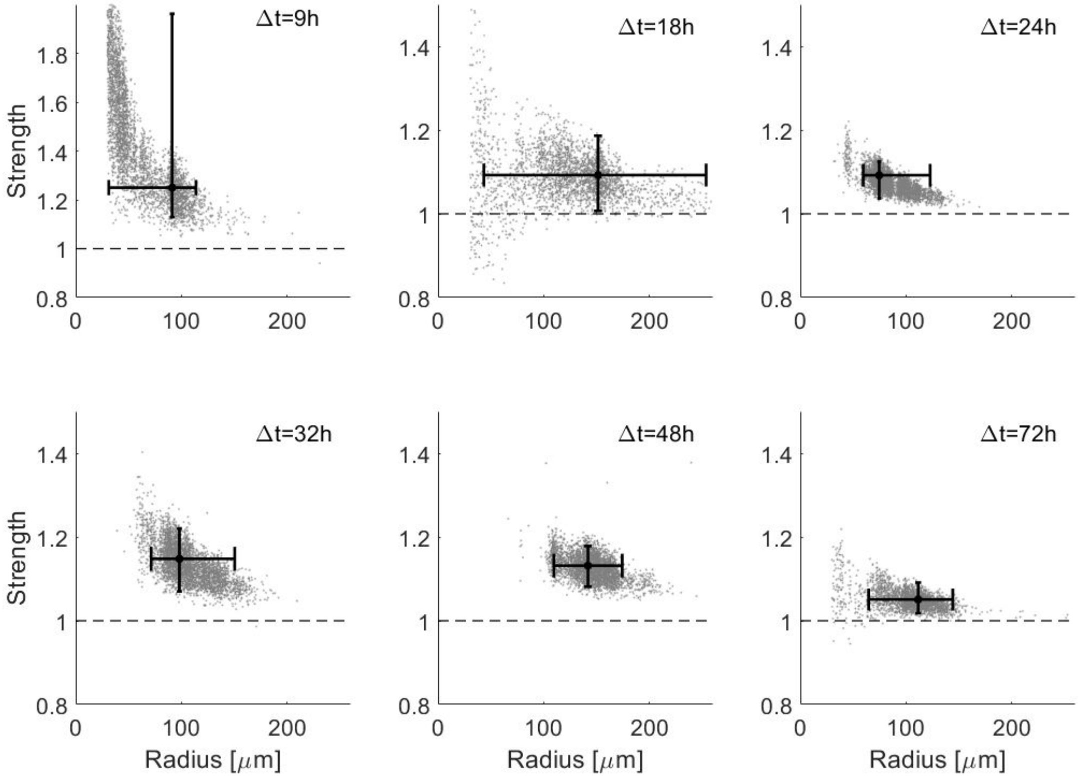


**Supplementary Figure 3.**
Parameter inference on all labelling intervals Δt. Most likely influence strength and radius for all hemispheres per labelling interval determined via posterior sampling density. Whiskers cover the 95% confidence intervals for radius and strength.

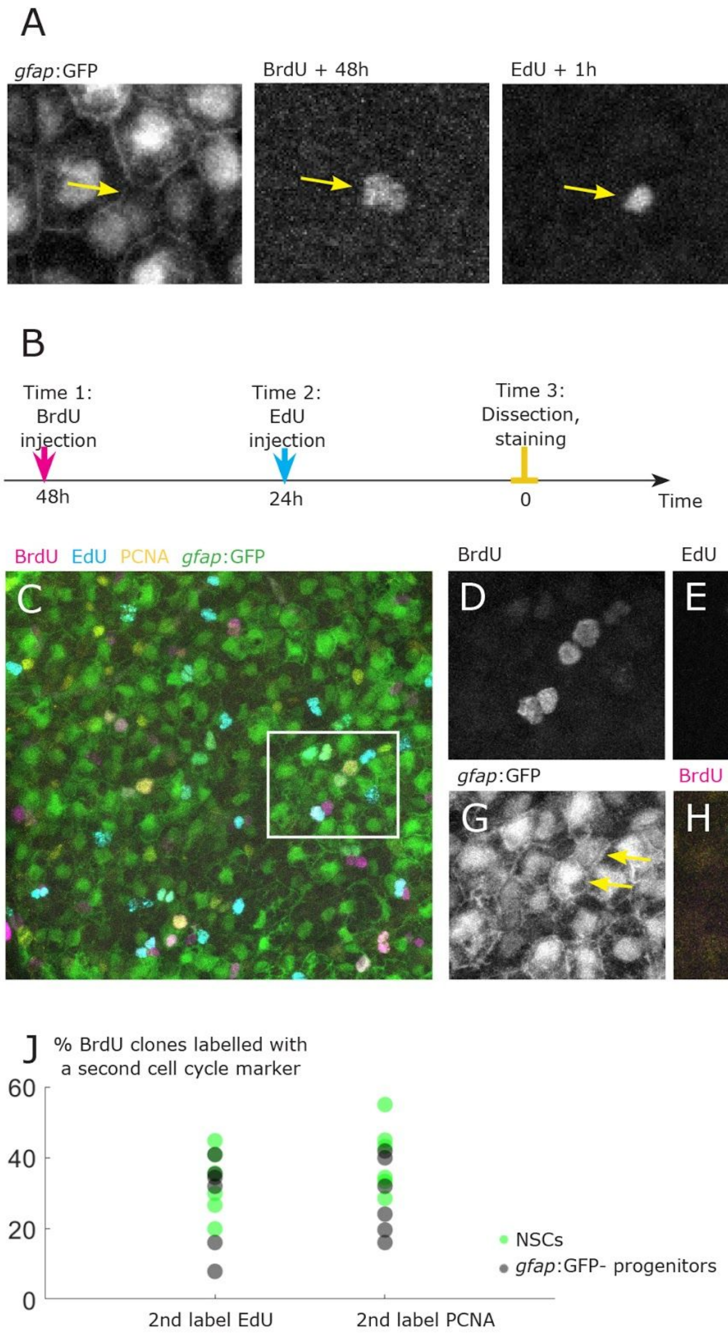


**Supplementary Figure 4.**
(A) Exemplary re-division of a *gfap*:GFP- progenitor doublet: The two close-by sister cells have divided 48h back (indicated by BrdU+), and the right sister is re-dividing (indicated by EdU+).
(B) Experimental setup for observing divisions at three time points within 2 days.
(C-F) Re-divisions marked with three cell cycle labellings at three time points: 2 days earlier by BrdU (D), one day earlier by EdU (E) and at the time of fixation by PCNA (F).
(G) Re-division of a *gfap*:GFP+ daughter cell pair (yellow arrows).
(H,I) Overlays.
(J) Re-divisions can be observed after 1 day (BrdU+EdU+) or after 2 days (BrdU+PCNA+), and have been quantified in 6 hemispheres from 4 brains. Higher percentages of re-divisions are observed compared to the experiments with a short pulse of EdU labelling at time 2 (injection 1h before killing, see Figure 4), probably due to the fact that more S-phases entries can be labelled with a longer duration of EdU availability in this experimental setup. Scale bars: 10μm.

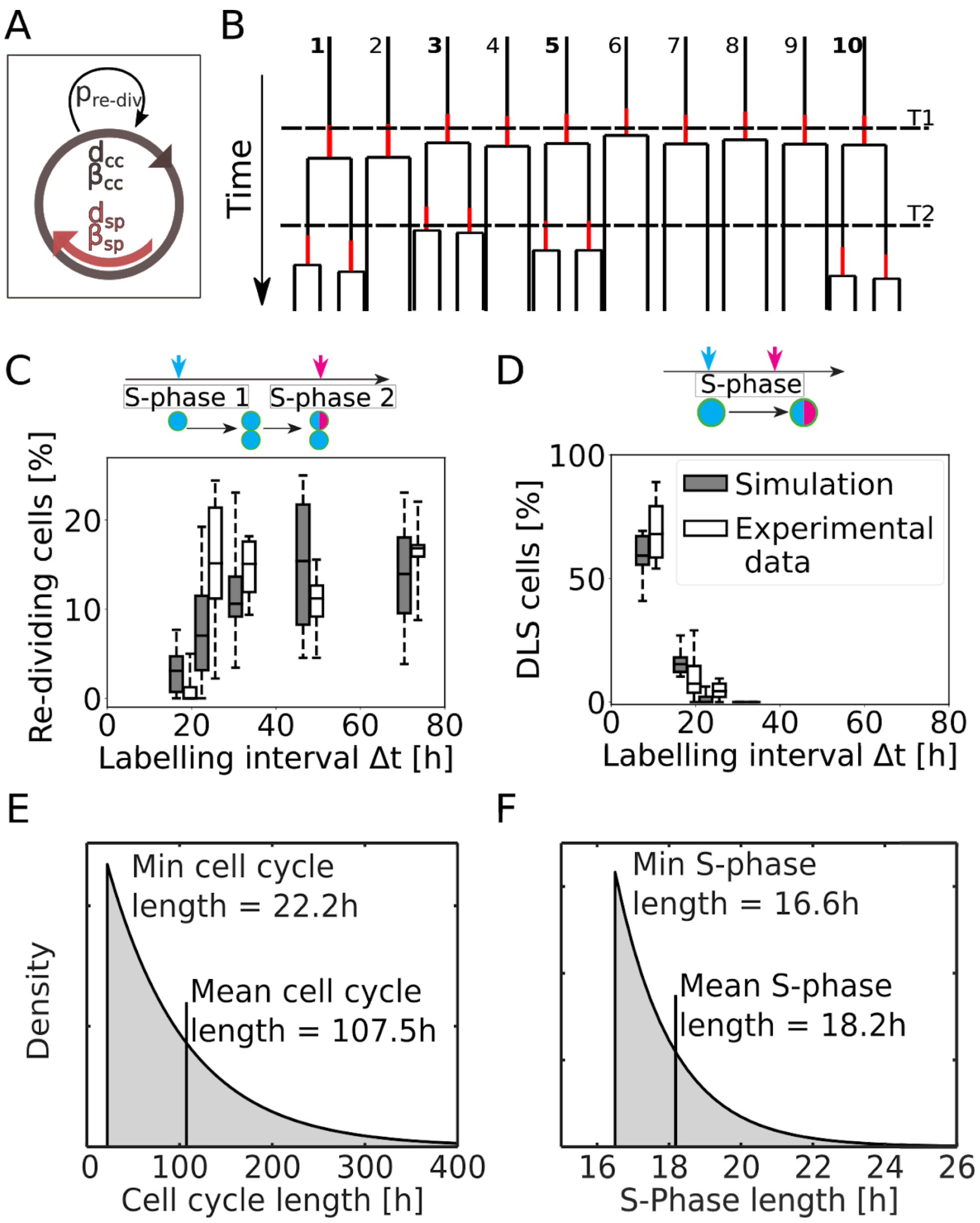


**Supplementary Fig 5.**

(A) We devise a non-spatial model of dividing NSCs with 5 parameters: the minimal cell cycle length $d_{cc}$ and minimal S-phase length $d_{sp}$ and their variability $\beta_{cc}$ and $\beta_{sp}$ respectively, parametrizing a lag-exponential distribution and the re-division probability $p_{re\text{-}div}$.

(B) We fit model parameters to observed re-division proportions and the percentage of DLS cells (S-phases labelled in red) by taking simulated measurements at T1 and T2. We observe a discrepancy between the real ($p_{re\text{-}div}$=0.38) and observed re-divisions frequency (0.15): From ten simulated cells dividing at T1, four cells (cell 1,3,5,10, shown in bold) re-divide. This corresponds to the 40%, which is close to $p_{re\text{-}div}$. However, our measurement at T2 only picks up two cells (cell 3 and 5) corresponding to only 20%, in line with the observed re-division frequency.

(C,D) Our re-division model fits the proportion of re-dividing and DLS cells for different labelling intervals.

(E) We find an extremely variable delayed exponential distributed cell cycle with a minimal length ($d_{cc}$) of 22h and $\beta_{cc}$ (=mean) of 85h while (F) S-phase length is tighter controlled with 16.6h minimal length ($d_{sp}$) and $\beta_{sp}$=1.5h.

**Supplementary Table 1.**
For each brain hemisphere (Experiment) with a particular labelling interval, we detail the number of NSCs, number of S-phases at labelling time 1 (T1) and labelling time 2 (T2), number of NSCs in S-phase at T1 and T2, number of re-divisions, number of NSC re-divisions, number of double labelled S-phases (DLS) and number of DLS in NSCs.

| Experiment | NSCs | T1 | T2 | T1 NSCs | T2 NSCs | Re-divisions | NSC re-divisions | DLS cells | DLS NSCs |
|---|---|---|---|---|---|---|---|---|---|
| Interval 9h H1 | 2102 | 79 | 57 | 44 | 28 | 0 | 0 | 47 | 25 |
| Interval 9h H2 | 2382 | 81 | 70 | 58 | 49 | 0 | 0 | 50 | 37 |
| Interval 9h H3 | 2351 | 61 | 64 | 40 | 46 | 0 | 0 | 35 | 24 |
| Interval 9h H4 | 2467 | 85 | 66 | 52 | 37 | 0 | 0 | 41 | 28 |
| Interval 18h H1 | 2867 | 103 | 53 | 65 | 17 | 2 | 0 | 3 | 0 |
| Interval 18h H2 | 2125 | 59 | 66 | 40 | 38 | 2 | 2 | 3 | 2 |
| Interval 18h H3 | 2389 | 86 | 96 | 41 | 41 | 0 | 0 | 6 | 4 |
| Interval 18h H4 | 1545 | 76 | 74 | 28 | 30 | 0 | 0 | 26 | 8 |
| Interval 24h H1 | 2524 | 111 | 132 | 59 | 66 | 0 | 0 | 0 | 0 |
| Interval 24h H2 | 2156 | 112 | 107 | 54 | 56 | 0 | 0 | 0 | 0 |
| Interval 24h H3 | 3304 | 120 | 111 | 63 | 46 | 0 | 0 | 0 | 0 |
| Interval 24h H4 | 3149 | 145 | 124 | 78 | 56 | 0 | 0 | 0 | 0 |
| Interval 24h H5 | 1565 | 91 | 123 | 55 | 60 | 6 | 5 | 1 | 1 |
| Interval 24h H6 | 1567 | 75 | 100 | 45 | 45 | 2 | 1 | 3 | 2 |
| Interval 24h H7 | 2097 | 28 | 101 | 14 | 44 | 0 | 0 | 0 | 0 |
| Interval 24h H8 | 2576 | 78 | 65 | 45 | 44 | 12 | 11 | 4 | 3 |
| Interval 24h H9 | 2099 | 82 | 44 | 35 | 28 | 10 | 9 | 3 | 0 |
| Interval 24h H10 | 2500 | 202 | 145 | 96 | 77 | 42 | 13 | 9 | 8 |
| Interval 24h H11 | 2859 | 294 | 202 | 117 | 117 | 63 | 28 | 7 | 5 |
| Interval 24h H12 | 2447 | 162 | 113 | 59 | 56 | 26 | 19 | 2 | 0 |
| Interval 24h H13 | 2305 | 161 | 77 | 57 | 46 | 22 | 17 | 2 | 1 |
| Interval 24h H14 | 1976 | 164 | 117 | 73 | 51 | 22 | 15 | 6 | 6 |
| Interval 24h H15 | 2549 | 202 | 146 | 68 | 58 | 36 | 12 | 7 | 3 |
| Interval 32h H1 | 2628 | 103 | 72 | 71 | 43 | 16 | 14 | 0 | 0 |
| Interval 32h H2 | 2542 | 82 | 67 | 46 | 31 | 8 | 5 | 0 | 0 |
| Interval 32h H3 | 2325 | 57 | 57 | 22 | 23 | 9 | 4 | 0 | 0 |
| Interval 32h H4 | 2801 | 155 | 73 | 63 | 28 | 16 | 5 | 0 | 0 |
| Interval 48h H1 | 1806 | 75 | 74 | 30 | 48 | 11 | 5 | 0 | 0 |
| Interval 48h H2 | 2243 | 111 | 73 | 49 | 43 | 15 | 11 | 0 | 0 |
| Interval 48h H3 | 3135 | 156 | 101 | 83 | 75 | 17 | 15 | 0 | 0 |
| Interval 48h H4 | 1882 | 56 | 67 | 23 | 29 | 7 | 2 | 0 | 0 |
| Interval 72h H1 | 2943 | 159 | 114 | 64 | 50 | 26 | 11 | 0 | 0 |
| Interval 72h H2 | 2260 | 95 | 69 | 44 | 28 | 14 | 7 | 0 | 0 |
| Interval 72h H3 | 3009 | 166 | 91 | 93 | 55 | 22 | 16 | 0 | 0 |
| Interval 72h H4 | 3319 | 169 | 129 | 88 | 79 | 24 | 14 | 0 | 0 |
| Interval 72h H5 | 3013 | 163 | 67 | 90 | 29 | 16 | 7 | 0 | 0 |
